# Stochasticity explains differences in the number of *de novo* mutations between families

**DOI:** 10.1101/2020.09.18.303727

**Authors:** Jakob M. Goldmann, Juliet Hampstead, Wendy S.W. Wong, Amy B. Wilfert, Tychele Turner, Marianne A. Jonker, Raphael Bernier, Martijn A. Huynen, Evan E. Eichler, Joris A. Veltman, George L. Maxwell, Christian Gilissen

## Abstract

The number of de novo mutations (DNMs) in the human germline is correlated with parental age at conception, but this explains only part of the observed variation. We investigated whether there is a family-specific contribution to the number of DNMs in offspring. The analysis of DNMs in 111 dizygotic twin pairs did not identify a significant family-specific contribution. This result was corroborated by comparing DNMs of 1669 siblings to those of age-matched unrelated offspring. In addition, by modeling DNM data from 1714 multi-offspring families we estimated that the family specific contribution explains approximately 5.2% of the variation in DNM number. Furthermore, we found no significant difference between the observed number of DNMs and those based on a stochastic Poisson process. We conclude that a family-specific contribution to DNMs is small and that stochasticity explains a large proportion of variation in DNM counts.

## Introduction

DNMs are drivers of genetic diversity and evolution and can also cause severe disease, such as intellectual disability, autism and schizophrenia (Veltman & Brunner, 2012). The number of single nucleotide DNMs per individual genome is ranges between 30 – 80 (Gillisen C *et. al*., 2014) and is correlated with the age of the parents at conception [Kong, Wong, Goldmann, Jonsson]. Ageing of fathers adds one DNM per year, while aging of mothers adds one DNM every four years. However, these two factors explain only part of the variation in DNM count, raising the question whether there are other factors aside from parental age that can be linked to the number of DNMs. Such factors could be endogenous, such as genetic variation in genes of DNA repair pathways (Goldberg ME *et. al*., 2019) or could be of external origin like ionizing radiation (Adewoye AB *et. al*., 2015; Holtgrewe M *et. al*., 2018) and environmental pollutants (Beal MA *et. al*., 2019; Ton ND *et. al*., 2018). Studies of multi-offspring families have suggested that the paternal age effect may differ significantly between families, where the mean yearly increase in DNMs per offspring with age of the fathers can vary from 0.2 to 3.2 DNMs per year (Rahbari R *et. al*., 2016; Sasani TA *et. al*., 2019).

Here, we analyzed DNMs from families with several offspring in order to investigate the family specific contribution to the variability in DNMs. We collected four cohorts from published whole-genome sequencing studies of families with multiple offspring, totaling 111 dizygotic twin pairs, 1714 multi-offspring families and 45 large families (median of 10 offspring) (**Table 1**). Because these cohorts were based on different sequencing and analysis methods they were analyzed separately after QC and used as independent replicates within this study (**Table 1, Supplementary Methods**).

**Table 1:**
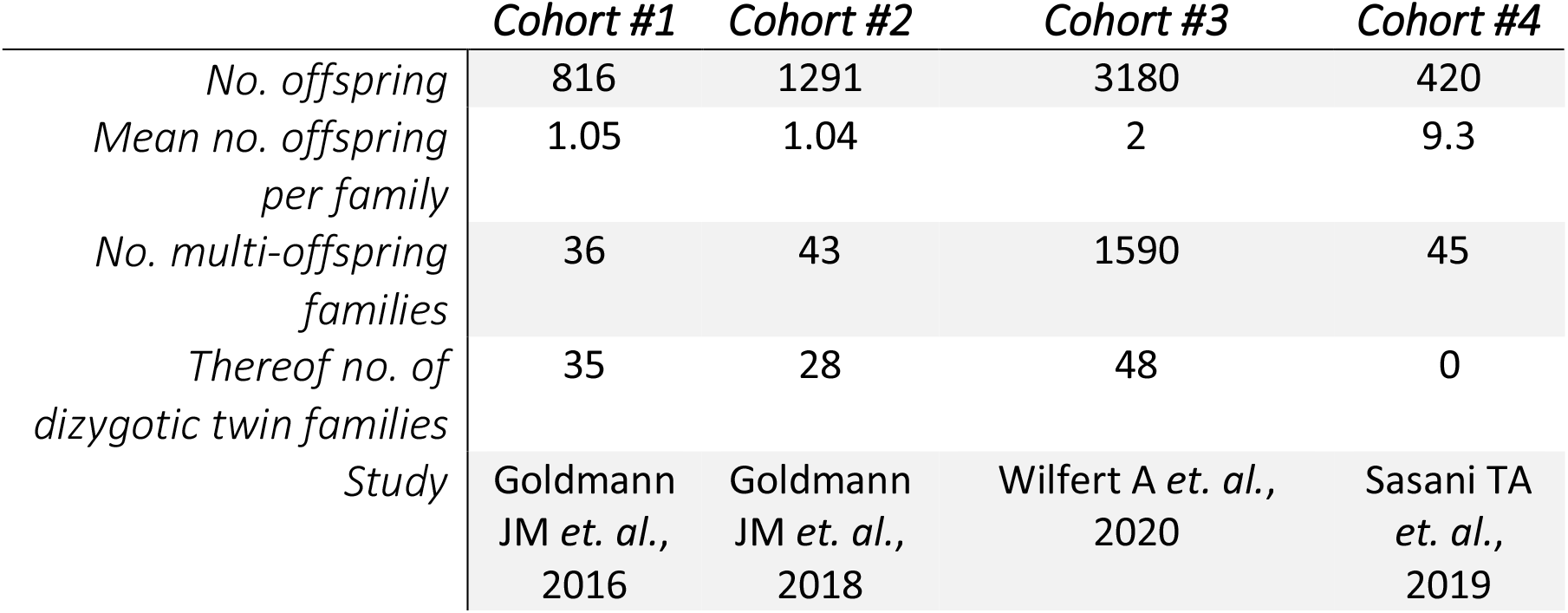
*Cohort descriptions*. Size of the cohorts of this study.

|  | *Cohort #1* | *Cohort #2* | *Cohort #3* | *Cohort #4* |
| --- | --- | --- | --- | --- |
| *No. offspring* | 816 | 1291 | 3180 | 420 |
| *Mean no. offspring per family* | 1.05 | 1.04 | 2 | 9.3 |
| *No. multi-offspring families* | 36 | 43 | 1590 | 45 |
| *Thereof no. of dizygotic twin families* | 35 | 28 | 48 | 0 |
| *Study* | Goldmann JM *et. al.*, 2016 | Goldmann JM *et. al.*, 2018 | Wilfert A *et. al.*, 2020 | Sasani TA *et. al.*, 2019 |

## Results

### Twins do not have a significantly different number of DNMs than age-matched unrelated children

If family-specific effects exist, this would cause unrelated individuals to have larger differences in the number of DNMs than siblings of a single family. The age at conception of father and mother are established factors that affect the number of DNMs in the offspring and need to be considered when comparing DNM counts between families. However, for dizygotic twins, we can directly compare the number of DNMs without correcting for parental age. The median differences in the number of DNMs for the dizygotic twins are 8, 8.5 and 9 for cohorts #1-3 respectively (**Figure 1a**; there are no twin families in cohort #4). The individual differences range from 0 DNMs to 29 DNMs. We did not observe significant trends in these differences nor a change in their variation with the age of the father (p-value for linear slope being different from zero p=0.31; Breusch-Pagan test for heteroscedasticity p=0.54, **Supplementary Note 1**).

**Figure 1:**
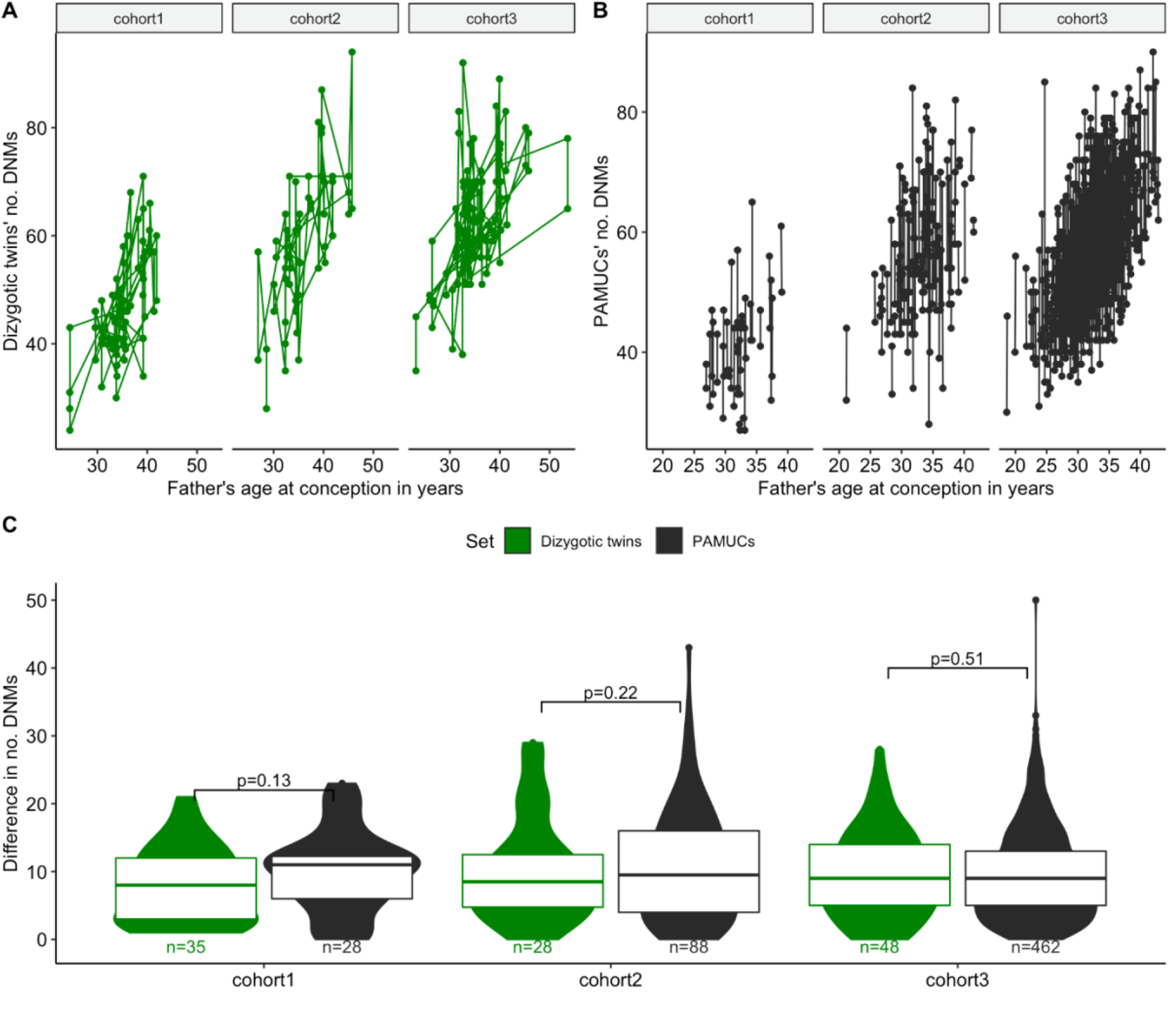
Dizygotic twins versus parental age-matched unrelated children. A: Numbers of DNMs of dizygotic twins in relation to age of the father. Twins are linked by lines. B: Numbers of DNMs of parental age matched unrelated controls (PAMUCs). C: Absolute differences in the number of DNMs of twins and PAMUCs. Numbers indicate sizes of sets, boxes indicate interquartile range and bold line indicates median.

We compared the differences in DNMs between twin pairs to those of 601 pairs of parental age-matched unrelated children (PAMUCs; **Methods**), and observed median differences of 11, 9.5 and 9 DNMs between the PAMUCs of cohorts #1-3 (**Figure 1b**). We did not detect a significant difference between the number of DNMs in twins and PAMUCs within any of the cohorts or for the combined dataset (**Figure 1c**, p=0.07; p=0.35; p=0.54 for cohorts #1-3 respectively; p-value for all datasets combined p=0.20, Wilcoxon Rank Sum test).

### Regression modelling of a larger cohort of unrelated children suggests that any family-specific contribution to de novo mutation rate must be small

The absence of a significant difference suggests that family-specific contributions to the number of DNMs may be small, or may be due to a lack of statistical power because of the relatively small number of dizygotic twins and PAMUCs (**Supplementary Note 2**). In order to increase our statistical power we included siblings of different ages and fitted a linear regression model that accounts for the parental ages at conception (**Supplementary Table 1**). In case of large family-specific effects, the difference in the observed number of DNMs between siblings should be significantly smaller compared to randomly matched pairs of offspring after accounting for parental age by our regression model. We compared the residual differences of families with two offspring (cohorts #1-3: 37, 42, 1590 sibling pairs) to unrelated other children in the same cohort but found no significant differences in any of the cohorts (**Figure 2**, cohorts #1-3: p=0.56; p=0.38; p=0.055; Wilcoxon Rank Sum test; there are no families with two offspring in cohort #4). These results suggest that any family-specific effect on the number of DNMs can only be small.

**Figure 2:**
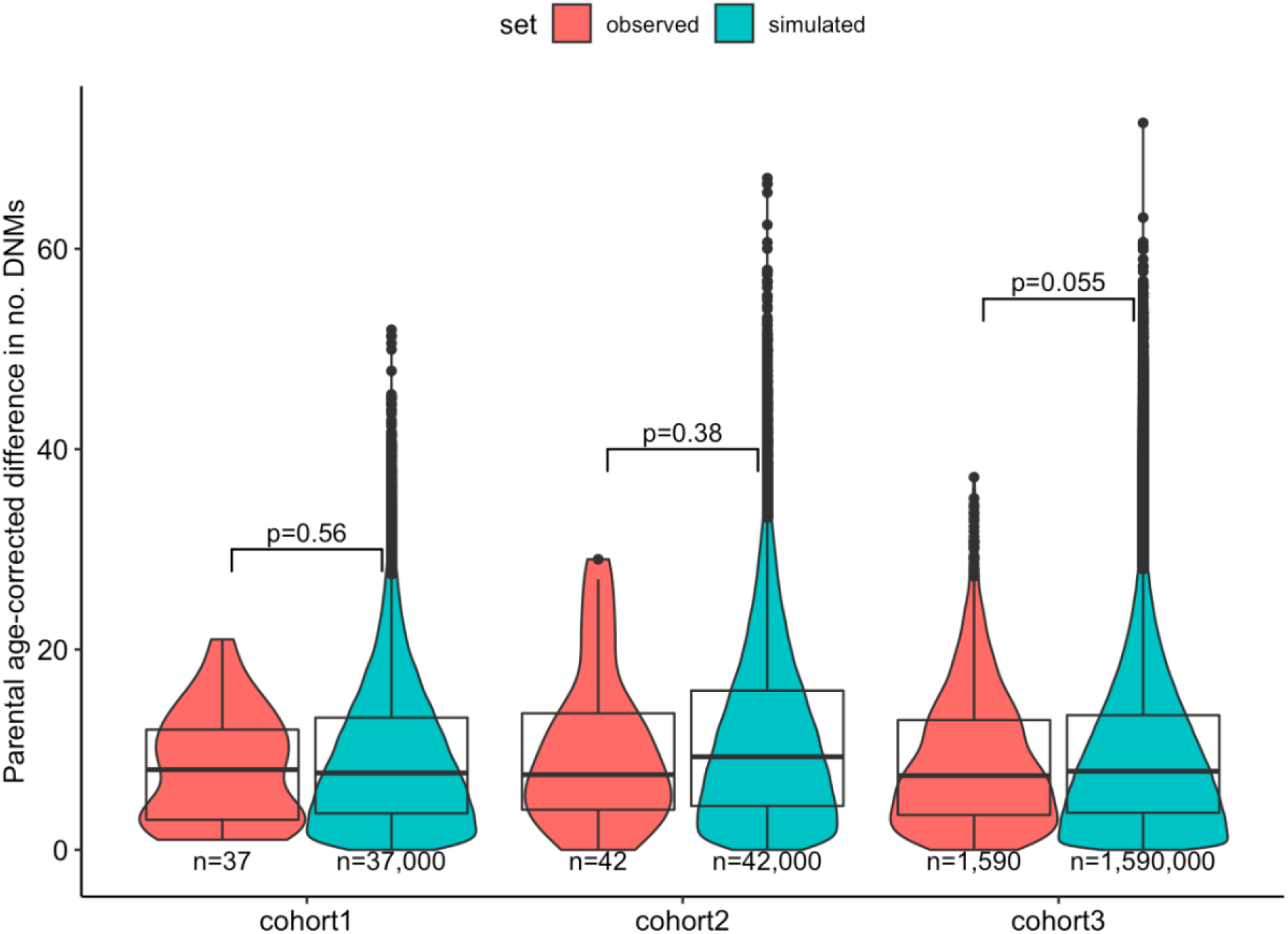
Absolute differences in DNM counts between siblings, corrected for parental age. Absolute differences of siblings were compared to differences of random combination of children. Numbers indicate sizes of sets, boxes indicate interquartile range and bold line indicates median.

### Random effects modelling allows direct estimation of the family-specific variance

We used a random effects model to directly estimate the potential impact of family-specific effects on the variation of DNM count. While the effects on the number of DNMs for paternal age and maternal age are fixed, we allow each family affiliation to add a specific number of DNMs to the total (**Methods**). We applied this model to our largest cohort with siblings (cohort #3), and the cohort of large families (cohort #4). We found that the point estimates for the variance components vary from 5.4% to 3.8% (**Figure 3, Supplementary Table 2, Supplementary Figure 2**). The mean of these familial variance component estimates, weighted by the number of offspring is 5.2%. This shows that family-specific effects can have only a very minor impact on *de novo* mutation rate.

**Figure 3:**
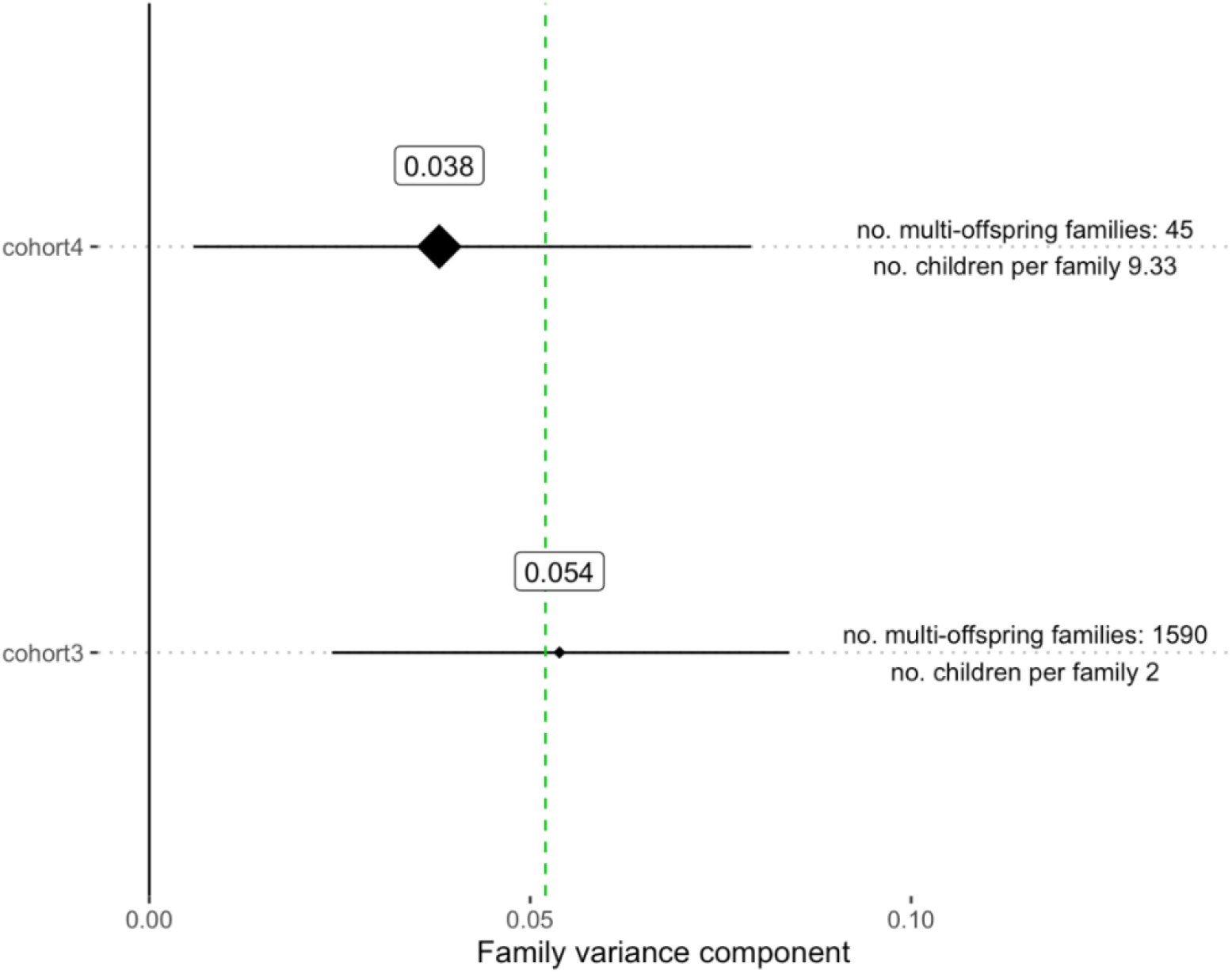
Family variance components. The error bars denote the 95% confidence intervals. The diamonds indicating the estimates are scaled according to the mean number of children per family. The vertical green line indicates the mean weighted by the number of multi-offspring families.

### Differences in DNM number between families can be simulated by a Poisson distribution

After having assessed the possibility of differences in DNM counts between families, we aimed to further understand the variance in DNM counts after parental age correction. We simulated mutation counts as a Poisson-distributed variable by first fitting a linear Poisson regression model to the observed DNM counts for obtaining the expected number of mutations dependent on parental age. For each family in a dataset, we obtained probabilities for all relevant mutation numbers and summed them. The resulting distributions closely resemble the observed DNM counts, with no significant differences detected in either the median nor variance (**Figure 4, Supplementary Table 3,** all Bonferroni-corrected p-values > 0.2). Additionally, when base pair changes are differentiated (C > A, C > G, C > T (non-CpG), CpG > TpG, T > A, T > C and T > G), we do not find significant differences between the observations and Poisson predictions, providing further support that family-specific effects may only contribute very little to the variability in the number of DNMs (**Supplementary Table 4, Supplementary Figure 1, Supplementary Note,** all Bonferroni-corrected p > 0.99).

**Figure 4:**
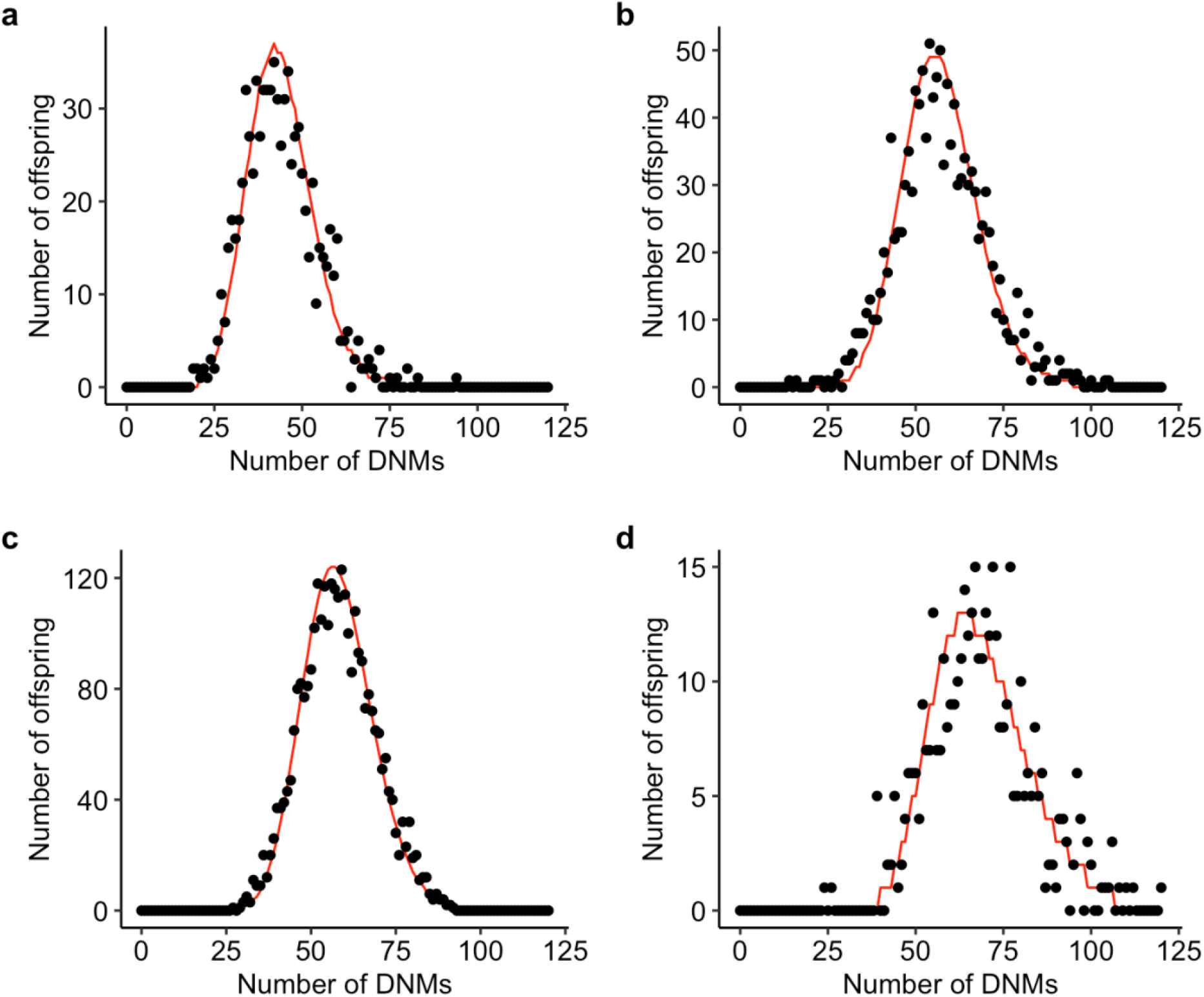
Modelling DNM as family-independent Poisson process. a,b,c,d: Simulations from cohorts #1-#4, respectively. Red lines depict Poisson-based predictions, black dots denote observations. **Supplementary Table 3** lists p-values for various test comparing predictions to observations.

## Discussion

Previously, Sasani *et al*. reported significant differences in the yearly increase rate of DNMs per family (Sasani TA *et. al*., 2019). Here, rather than focusing on the yearly increase we studied the total number of DNMs, and show that differences between families are very small. Thus, while families might vary in mutation accumulation rate, the number of DNMs passed on to offspring appears relatively constant after parental age correction. Our results indicate that there is no overall large family-specific effect in the general, healthy population (the upper border of 95% confidence intervals is 8.4% of the total variance in DNM numbers). However, our analyses do not exclude rare individual families in which genetic or environmental factors have given rise to a significantly increased DNM rate.

Our finding that stochasticity appears to play a major role in the accumulation of mutations in germline cells is of potential interest for reproductive medicine. It implies that even among the germline cells of older males, which on average have accumulated many mutations, there still may be lineages of cells with relatively few mutations. If it would be possible to identify these low-mutation load spermatogonia lines and isolate the corresponding sperm, the risk for giving birth to genetically diseased offspring could be lowered.

## Methods

### Cohorts

Cohort #1 is the Inova Translational Medicine Institute (ITMI) Premature Birth Study cohort with 816 healthy newborns being born at the Inova hospital (Goldmann JM *et. al*., 2016). One third of probands (219) was born prematurely (gestational age < 37 weeks). Cohort #2 is the ITMI Childhood Longitudinal Cohort Study cohort (Goldmann JM *et. al*., 2018). Cohort #3 is a combination of the SSC, TASC, and SAGE study cohorts and were sequenced at the New York Genome Center (Wilfert A *et. al*., 2020). Cohort #4 is a collection of large families from the Centre d’Etude du Polymorphisme Humain (CEPH) consortium (Sasani TA *et. al*., 2019; Dausset J *et. al*., 2020). The cohorts are compared in **Supplementary Table 1**. Specifics of the custom pipelines used for DNM calling in each cohort are available in the appropriate references. Where the age at conception was not available, we used the age at birth accordingly.

In all three cohorts, parental age-matched unrelated children (PAMUCs) were identified by scanning for pairs of children where the sum of the differences in parental ages was less than 43 days.

### Analysis of parental-age corrected DNM counts

We fitted a linear model predicting the number of DNMs based on the age of mother and father at conception (**Supplementary Figure 3**). For each offspring, we calculated the residuals of the model by subtracting the observed DNM count from the prediction. For each family with two offspring, we calculated the absolute difference of the two offspring’s residuals. We compared these parental age-corrected difference in DNMs to the differences of two offspring randomly sampled from the same cohort. We re-sampled the family labels 1.000 times. We used Wilcoxon Rank Sum tests for assessing statistical significance.

In cohort 3, each family contained one patient with an autism-spectrum disorder and one unaffected sibling. We could not detect a significant difference in the parental-age corrected number of de novo mutations between these two groups (**Supplementary Figure 4**).

### Estimating the variance component of familial influences

We model the number of DNMs of an individual as the sum of a baseline expectation, the paternal age effect and the maternal age effect and a residual error term. More specifically, the number of DNMs *X* of an individual *i* is

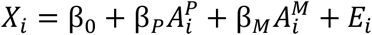

where β_*o*_ is the baseline number of DNMs that occur during prenatal development, and β_P_ and β_M_ are the strengths of paternal and maternal age effects, respectively, supplied in DNMs per year. The factors 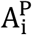 and 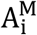 are the ages of father (paternal) and mother (maternal) of the respective individual at conception. The residual error is captured by the random effects term E_i_ that is specific to every individual.

To allow for possible familial influences on the number of DNMs, we added a familial influence factor *F_j_*, which is a random effects term specific to every family *j*.

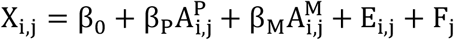

The introduction of this term allows us to estimate the variance introduced to the model by family-specific influences. For this, the model is fitted to observed data using the R statistical environment with the package “lmer” for fitting the linear models with random effects (Bates D *et. al*., 2015). We obtained the variance components of all factors in the model with 95% confidence intervals by applying the function “rpt” R package “rptR” with 500 fold bootstrapping, which estimates variance components for both fixed and random effects (Stoffel MA *et. al*., 2017).

### Batch effect estimation

We model the batch effect in the same way as the family effect, that is by including a batch-specific random effects term to the regression formula. Fitting this term to every batch allowed for estimation of inter-batch variation.

Nevertheless, this approach requires batch annotation to be both present and sufficiently disjunct from the family annotations, such that the fitting algorithm can robustly differentiate both effects. For cohorts #1 and #3, such annotations were available to us; these were version numbers of the software pipeline for cohort #1 and the date of the sequencing run for cohort #3.

### Poisson simulations of DNM counts

We predicted the number of DNMs for every offspring by fitting a Poisson regression model based on parental ages. We use the “dpois”-function of the R statistical environment for predicting the density of a Poisson distribution with the predicted number of DNMs as expectation value. We obtained predictions for each number of DNMs from 0 to 75. Across each cohort, we summed up these densities for all offspring and rounded the results.

To compare predicted densities to observed values, we used two sets of statistical tests. First, we used a Wilcoxon rank-sum test to assess for differences in the median of the distributions. Second, we used a group of tests to assess differences in the variance of the distributions, including Levene’s Test and the Fligner-Killeen Test for heterogeneity of variance, the Ansari-Bradley Test and Mood’s Test for the difference in scale parameters as well as the parametric F-Test for comparison of variances.

### Multiple testing correction

P-values were corrected for multiple testing by Bonferroni’s method where indicated.

## Data Access

Data from all 4 cohorts used in this study and code to reproduce analysis and figures will be made available on GitHub. DNM data from each of the four cohorts (#1-#4) is available in the supplementary materials of the publication indicated in **Table 1** (Goldmann JM *et. al*., 2016; Goldmann JM *et. al*., 2018; Wilfert A *et. al*., 2020; Sasani TA *et. al*., 2019).

## Acknowledgements

The authors declare no acknowledgments.

## Author Contributions

JMG, CG and JAV conceived and planned the study. JMG and MAJ carried out the analysis. JMG, JH, WSWW, TT, MAJ, MAH, EEE, JAV, CG interpreted the results. JMG, WSWW, ABW, TT, RB, EEE, GLM acquired the data. JMG, JH, WSWW, MAH, JAV, CG drafted the manuscript. All authors contributed to the final version of the paper. Correspondence and material requests can be addressed to CG.

## Disclosure Declaration

The authors declare no competing interests.

